# Ammonium retention by Amberlite IRC-748 resin: useful for concentration assessments

**DOI:** 10.64898/2026.05.05.722854

**Authors:** Haikuo Zhang, Harald Neidhardt, Steffen Seitz, Thomas Scholten, Yvonne Oelmann

## Abstract

Chelating ion exchange resins are widely used to eliminate metal interferences in the analysis of ammonium (NH_4_^+^) in soil extraction solutions. However, their potential to co-adsorb NH_4_^+^ remains underexplored. Here, synthetic metal ion solutions containing 6–30 mg L^-1^ NH_4_^+^ and the metal cations Ca^2+^, Mg^2+^, Cu^2+^, Mn^2+^, and Zn^2+^ were treated with Amberlite IRC-748 resin. The resin efficiently removed Ca^2+^ (−42.2%), Mg^2+^ (−21.1%), Cu^2+^ (−99.9%), Mn^2+^ (−56.9%), and Zn^2+^ (−93.6%). However, NH_4_^+^ losses of 2.2–5.6% were observed, indicating concentration-dependent co-adsorption. While these losses may be acceptable for concentration measurements via routine assays such as photometric analysis, they may still affect the accuracy of high-precision N analyses that rely on quantitative NH_4_^+^ recovery. This highlights a methodological caveat for resin-treated samples, especially in low-NH_4_^+^ environments. We therefore recommend including recovery assessments and correction factors when using chelating resins to improve accuracy in NH_4_^+^ quantification.

## Introduction

Quantifying ammonium (NH_4_^+^) in soil is critical for understanding nitrogen (N) dynamics, especially in fertilized or contaminated environments where metal interference complicates analysis (Mulvaney 1996). Divalent metal ions such as Ca^2+^, Mg^2+^, Cu^2+^, Mn^2+^, and Zn^2+^ can interfere with the indophenol blue reaction by complexing with reagents or inhibiting oxidation, thereby causing underestimation of NH_4_^+^ concentrations measured by UV-colorimetry (Willis et al. 1993; Mulvaney 1996). To overcome this limitation, chelating ion exchange resins such as Amberlite IRC-748 are widely used due to their strong affinity for divalent and transition metal ions (Mumford et al. 2008; Yu et al. 2015). These resins are commonly recommended in protocols for environmental soil extract preparation (Yu et al. 2015; Thinh et al. 2021; Chu et al. 2022). However, most existing studies have focused primarily on the efficiency of removing metals by resin, and their potential to inadvertently adsorb NH_4_^+^ remains largely unexamined (Lin and Juang 2007). Accurate NH_4_ ^+^ recovery is especially critical for high-precision N analyses that rely on quantitative NH_4_^+^ recovery, such as N isotope analysis, where even small losses may lead to substantial isotope fractionation, which biases results (Houlton et al. 2006; Craine et al. 2015).

Furthermore, any unaccounted loss of NH_4_^+^ during sample pre-treatment may lead to biased interpretations or flawed nutrient budgeting (Lehmann et al. 2001). The extent to which NH_4_^+^ is inadvertently adsorbed by the resin may depend on the initial NH_4_^+^ concentration in the extraction solutions (Keithley et al. 2021). Without proper quantification of this loss, studies that rely on resin-treated extracts may systematically underestimate the availability of soil NH_4_^+^ (Lehmann et al. 2001). To date, quantitative assessments of NH_4_^+^ loss during resin-based metal removal under different NH_4_^+^ concentrations and realistic ionic matrices remain scarce (Ding and Sartaj 2016).

To address these knowledge gaps, we conducted a controlled laboratory experiment using Amberlite IRC-748 resin to evaluate its potential to co-adsorb NH_4_^+^ during metal removal. A synthetic multi-cation solution was designed to mimic the ionic composition of fertilized and metal-contaminated soils. We quantified NH_4_^+^ losses across different initial concentrations and analyzed the resin’s selectivity toward major cations. By quantifying these effects, our study provides methodological insights into potential analytical biases in NH_4_ ^+^ determination following resin-based metal removal.

## Materials and methods

### Synthetic solution preparation

We prepared a synthetic solution containing 4 mM KCl, 4 mM CaCl_2_, 1 mM MgCl_2_, 0.5 mM CuCl_2_, 2 mM MnCl_2_, and 2 mM ZnCl_2_ to simulate fertilized and high-metal soil extracts. These metal ions were added simultaneously to approximate the cationic complexity of fertilized and metal-contaminated soils. The synthetic solution was also spiked with NH□□ standard solution at four concentration levels: 6, 15, 25, and 30 mg L^-1^, each with four replicates, and two parallel treatments (experimental and control treatments, see below) were designed.

### Resin preparation

Amberlite IRC-748 (Rohm and Haas, Philadelphia, USA) was pre-washed with 0.1 M HCl to remove residual metals, rinsed with deionized (DI) water, treated with 0.1 M NaOH to restore the Na^+^ form, and rinsed to near-neutral pH. Excess water was removed by vacuum filtration through a filter with a 0.45 μm membrane (Carl Roth, Karlsruhe, Germany), with the resin retained on the filter for subsequent use.

### Experimental treatment (adsorption with metal cations)

For each NH_4_ ^+^ level, the NH_4_ ^+^-containing solution was prepared using the synthetic solution as the base solution. Based on the total metal ion content and the reported adsorption capacity, subsamples of 20 mL were transferred to 50 mL polypropylene centrifuge tubes and mixed with 100 mg of the pre-treated resin. The resin dosage was chosen based on the total metal ion content in the 20 mL solution and the reported adsorption capacity of Amberlite IRC-748 resin for divalent metals. Tubes were shaken horizontally (150 rpm, 8 h, room temperature). After shaking, the solutions were analyzed for the concentrations of residual NH_4_ ^+^ and metal ions colorimetrically by the indophenol blue method at 660 nm using a continuous flow analyzer (CFA, Seal Analytical AA3, Hamburg, Germany) and an inductively coupled plasma optical emission spectrometry (ICP-OES, Jena Analytik, Jena, Germany), respectively.

### Control treatment (NH_4_^+^ in the demetallized background solution)

To ensure that the control and resin-treated samples shared an identical background matrix during NH_4_^+^ determination, the same synthetic solution was pre-treated with the resin under identical conditions (100 mg per 20 mL, 150 rpm, 8 h). This pre-treated solution retained the same residual metal background as the resin-treated samples in the experimental treatment. It was then spiked with NH_4_^+^ to four target concentrations, serving as the control treatment, and analyzed by CFA. This design eliminated matrix effects from residual metals after the resin-treated, ensuring that any differences in measured NH_4_^+^ concentrations reflected resin co-adsorption rather than analytical bias.

### Blank corrections

Blank corrections were performed using DI water alone and DI water with resin added. The corresponding background NH_4_ ^+^ concentrations were subtracted from those measured in the adsorption treatments to eliminate potential contributions from NH_4_^+^ present in the DI water or resin.

### Data analysis

We observed no detectable loss of solution volume during the adsorption experiments; therefore, the difference in NH _4_ ^+^ concentration between the control and experimental treatments was interpreted as NH_4_^+^ loss resulting from co-adsorption or competitive exchange with the resin. The concentration and percentage of NH_4_^+^ loss were computed as:

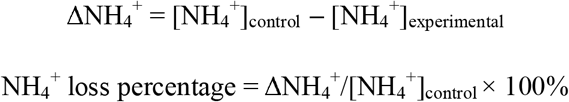

where the ΔNH_4_^+^ is the concentration difference of NH_4_^+^ between control and experimental treatments, the [NH_4_^+^]_control_ is the NH_4_^+^ concentration in the demetallized control treatment, and the [NH_4_ ^+^]_experimental_ is the NH_4_ ^+^ concentration in the experimental treatment.

Furthermore, for each cation (K^+^, Ca^2+^, Mg^2+^, Cu^2+^, Mn^2+^, Zn^2+^), the concentration and percentage of metal-ion removal by the resin in the experimental treatment were calculated as follows:

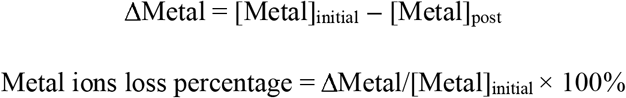

where ΔMetal is the concentration difference of metal ions before and after resin treatment, the [Metal]_initial_ is the concentration of metal ions before resin contact, and the [Metal]_post_ is the concentration of metal ions after resin treatment.

Data are reported as mean ± SD (n = 4). Differences in concentration between control and experimental treatments for each metal ion/NH _4_^+^ level, differences in loss percentage among metal ions, and among NH_4_^+^ concentration levels were assessed by one-way ANOVA followed by Tukey’s HSD for multiple comparisons (α = 0.05).

## Results & discussion

Our findings showed that treatment of simulated cationic solutions with Amberlite IRC-748 resin effectively removed most divalent and transition metal ions, but also resulted in losses of NH_4_^+^ across all tested concentrations (2.2–5.6%, Figs. 1 and 2). The ion-specific removal revealed distinct selectivity of the resin: while K^+^ concentrations remained unchanged after resin treatment, significant reductions were observed for Ca^2+^ (−42.2%, *p*<0.001), Mg^2+^ (−21.1%, *p*<0.001), Cu^2+^ (−99.9%, *p*< 0.001), Mn^2+^ (−56.9%, *p*<0.001), and Zn^2+^ (−93.6%, *p*<0.001) (Fig. 1). These findings align with the reported strong affinity of Amberlite IRC-748 for heavy metals and transition metal cations, particularly Cu^2+^ and Zn^2+^, which were nearly completely removed by the resin (Abusultan et al. 2023). Unchanged K^+^ concentration (0%, *p* > 0.05) suggests that monovalent ions are not efficiently retained by the resin under the tested conditions, as expected (Yu et al. 2009).

**Fig. 1.**
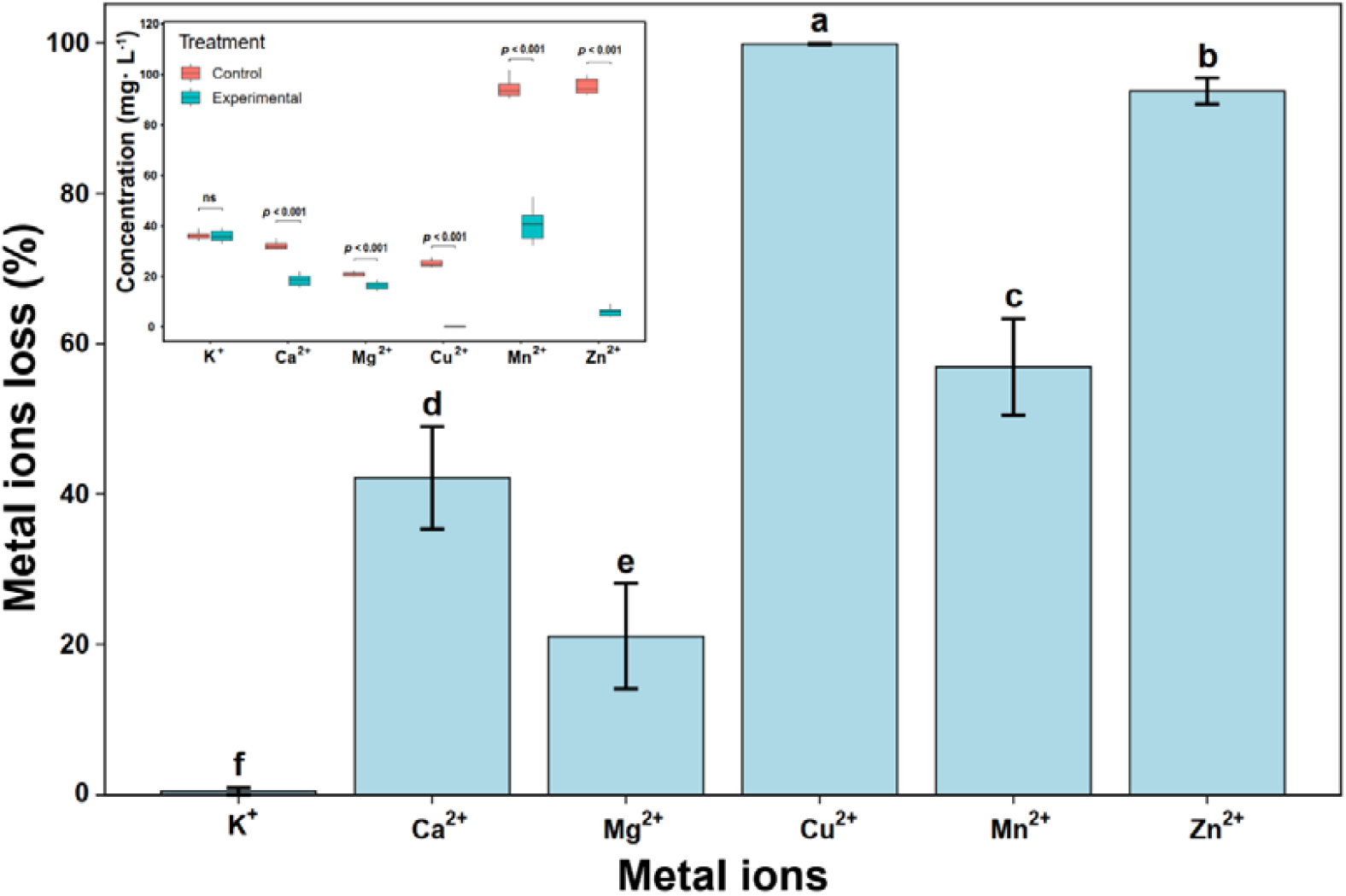
Metal ions loss percentage and concentrations comparison before and after resin treatment in 20 ml cationic solution. “ns” indicates no significant difference; The *p*-values and different lowercase letters indicate statistically significant differences based on one-way ANOVA followed by Tukey’s Honestly Significant Difference test.

**Fig. 2.**
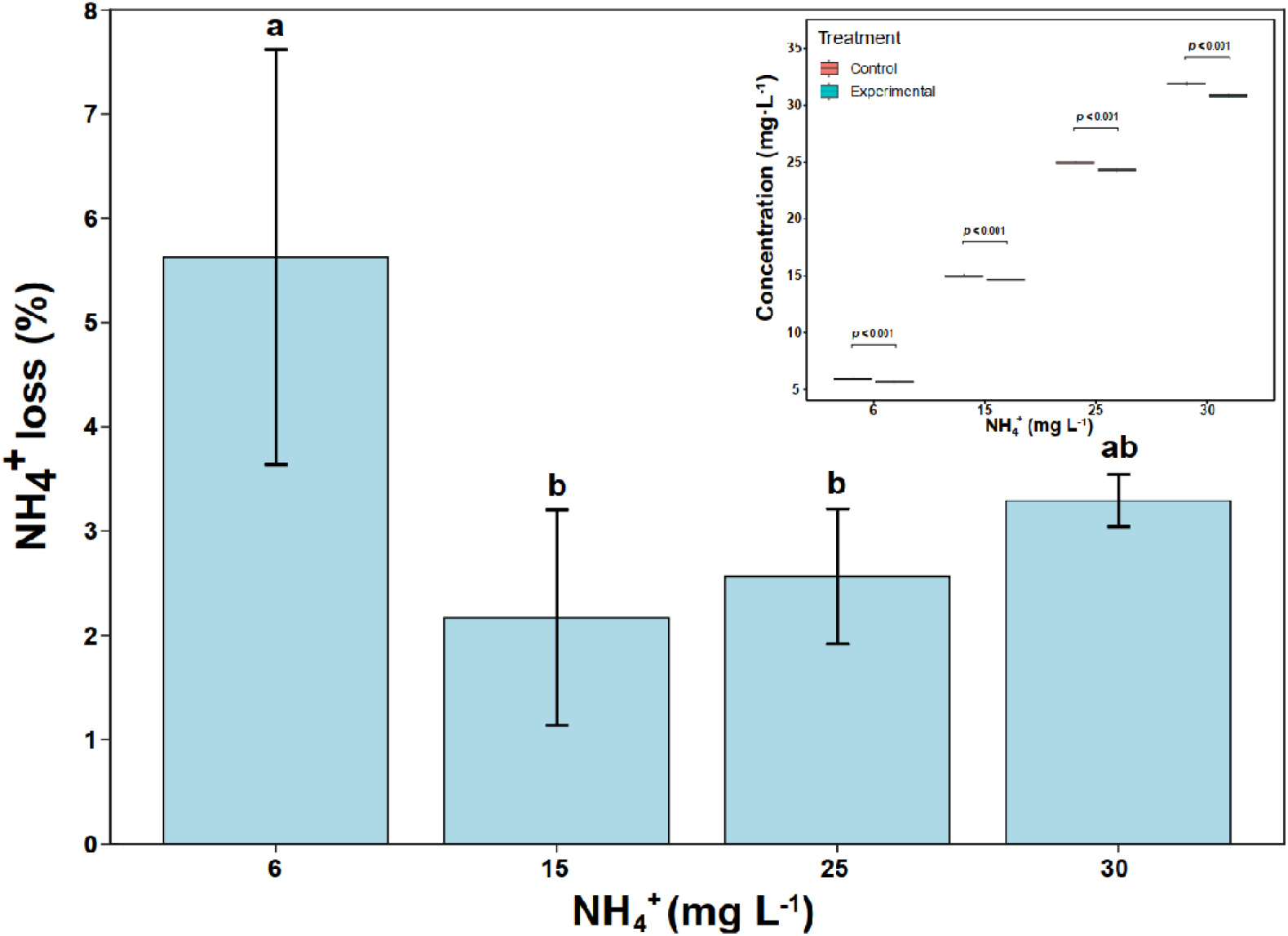
The NH_4_^+^ loss percentage and concentrations comparison before and after resin treatment in a 20 ml cationic solution. The different lowercase letters indicate statistically significant differences based on one-way ANOVA followed by Tukey’s Honestly Significant Difference test.

Compared with the control treatment, the NH_4_^+^ concentration in the experimental treatment was significantly reduced (Fig. 2). Specifically, the average NH _4_^+^ losses in the 20 ml reaction volume were 0.0068 mg, 0.0065 mg, 0.0128 mg, and 0.0210 mg in the 6, 15, 25, and 30 mg L^-1^ NH_4_^+^ concentration levels, respectively. Despite small absolute amounts, these losses represented 2.2% to 5.6% of the initial NH_4_^+^ added, depending on concentration. This resin−NH_4_^+^ interaction mechanism may arise from weak electrostatic or hydrogen-bonding interactions between NH_4_ ^+^ and the iminodiacetate (–N(CH_2_COOH)_2_) functional groups of the IRC-748 resin, or from physical entrapment of NH_4_^+^ within the resin’s porous structure, particularly at high ionic strength (Chen et al. 2015; Abusultan et al. 2023).

These findings highlight a key methodological consideration for studies involving NH_4_^+^ quantification in metal-rich soil extracts. Although the use of chelating resins, such as Amberlite IRC-748, is common in pre-treatment protocols to eliminate metal interferences in colorimetric and flow-based NH_4_^+^ assays, our results suggest that such pre-treatment may lead to an underestimation of NH_4_^+^ availability.

From a practical standpoint, our results do not preclude the use of resins like Amberlite IRC-748 in soil sample preparation, but suggest that analytical workflows should include quantification of potential NH_4_^+^ losses and possibilities for recovery. This could involve the inclusion of procedural blanks and correction factors when reporting NH_4_ ^+^ concentrations with resin treatment.

In conclusion, while Amberlite IRC-748 is effective in removing metal ions from complex aqueous matrices, it also causes a quantifiable bias in NH_4_ ^+^ concentration measurements due to co-sorption effects. Although the observed NH_4_^+^ losses were small (ranging from 2.2% to 5.6%), and within the acceptable range for concentration-based methods such as CFA, such losses are not acceptable for high-precision N analyses that rely on quantitative NH_4_^+^ recovery. Therefore, the use of chelating resins like Amberlite IRC-748 is not recommended for N isotope tracing studies involving NH_4_^+^. Our findings call for greater analytical scrutiny in NH_4_^+^ analysis of metal-rich soil extracts and highlight the need to correct for NH_4_^+^ adsorption when applying resin-based pre-treatment protocols in soil environmental studies.

## Acknowledgements

We would like to thank Ms. Rita Moegenburg and Ms. Sabine Flaiz from the Laboratory of Soil Science and Geoecology at the Institute of Geography, University of Tübingen, for their technical support with ICP-OES and CFA analysis, respectively.

## Data availability statement

The data that support the findings of this study are available from the corresponding author upon reasonable request.

